# Scaling of subcellular structures with cell length through decelerated growth

**DOI:** 10.1101/2021.03.01.433461

**Authors:** Shane G. McInally, Jane Kondev, Bruce L. Goode

**Affiliations:** Department of Biology, Brandeis University, Waltham, MA, 02454, USA; Department of Physics, Brandeis University, Waltham, MA, 02454, USA

## Abstract

How cells tune the size of their subcellular parts to scale with cell size is a fundamental question in cell biology. Until now, most studies on the size control of organelles and other subcellular structures have focused on scaling relationships with cell volume, which can be explained by limiting pool mechanisms. Here, we uncover a distinct scaling relationship with cell length rather than volume, revealed by mathematical modeling and quantitative imaging of yeast actin cables. The extension rate of cables decelerates as they approach the rear of the cell, until cable length matches cell length. Further, the deceleration rate scales with cell length. These observations reveal a new mode of scaling that senses the linear dimensions of a cell.

**One Sentence Summary:** As actin cables in yeast cells grow longer, their extension rate decelerates, enabling the cable length to match cell length.

## Main Text

The size of a cell’s internal parts are scaled to its overall size. This size-scaling behavior has been demonstrated for organelles as well as large macromolecular assemblies, illustrating the broad importance of adapting the size of internal structures to the geometric dimensions of the cell (*1–10*). A popular model of cellular scaling is the limiting pool mechanism, wherein maintaining a constant concentration of molecular components enables the subcellular structure to increase in size proportionally with cell volume (*11*). This allows larger cells to assemble larger structures, since the total number of molecular building blocks increases proportionally with cell volume. Additionally, this mechanism is biochemically simple because it does not require active processes that dynamically tune concentrations or activity levels of proteins involved in the construction (*12*). Indeed, cells appear to use a limiting pool mechanism to scale the size of their nucleoli, centrosomes, and mitotic spindles (*3–6*, *8*, *10*). However, limiting pool models cannot explain how the size of a linear subcellular structure scales with the linear dimensions of a cell, rather than its volume. Namely, these mechanisms predict that a two-fold increase in the radius of a spherical cell will increase the length of a linear structure eight-fold, following the eight-fold increase in cell volume. This suggests that other mechanisms must account for how some subcellular structures are scaled with the linear dimensions of a cell.

Polarized actin cables in *S. cerevisiae* are an example of a linear structure that appear to grow to match the linear dimensions of the cell in order to effectively deliver secretory vesicles (*13*). These cables are linear bundles of crosslinked actin filaments assembled by formins, which extend along the cell cortex and serve as tracks for intracellular transport of cargo from the mother cell to the growing bud, or daughter cell. Complementary sets of cables are assembled by two formins, Bni1 at the bud tip and Bnr1 at the bud neck (*14*). Throughout the cell cycle, cables are continuously polymerized, turn over at high rates, and appear to grow until they reach the back of the mother cell (*15–17*). This prompted us to more rigorously investigate the relationship between cable length and cell size.

We started by comparing cable lengths to the lengths of the mother cells in which they grew. Cables were imaged in fixed wildtype haploid cells using super-resolution microscopy. Cable lengths were measured from their site of assembly (the bud neck) to their distal tip in the mother cell (note that mother cell and cell are synonymous, and used interchangeably from this point). Average cable length and average cell length were remarkably similar (4.6 ± 0.3 μm and 4.5 ± 0.2 μm, respectively). Because the cables in fixed cells are at different stages of growth, their lengths vary considerably. Further, because cables grow along the cortex of an ellipsoid shaped cell, their length can exceed the length of the cell while not growing past the back of the cell. Therefore, a cable that grows from the bud neck to the back of the cell is expected to be slightly longer than the direct distance between these two points.

The observations above led us to ask whether the relationship between cable length and cell length is maintained as cell size increases. To address this, we compared cable lengths in haploid and diploid cells, and *cdc28-13*^*ts*^ temperature sensitive mutants that grow abnormally large. Diploid mother cells had an ~2-fold increase in volume compared to haploid mother cells (81.5 ± 6.2 μm^3^ and 44.6 ± 4.5 μm^3^, respectively) (Fig. 1, A, B, and E), consistent with previous studies (*18*). The *cdc28-13*^*ts*^ strain exhibited a normal haploid mother cell size at the permissive temperature, and ~5-fold increase in volume (199.3 ± 2.7 μm^3^ versus 40.7 ± 2.3 μm^3^) at the restrictive temperature (Fig. 1, C, D, and E; and fig. S1, A and B)(*19*). Accordingly, cell length increased with cell volume (Fig. 1F). Cable length was greater in diploids (6.2 ± 0.7 μm) compared to haploids (4.6 ± 0.3 μm), and greater in induced (8.2 ± 0.4 μm) compared to uninduced (4.4 ± 0.1 μm) *cdc28-13*^*ts*^ cells (Fig. 1G). However, the distribution of cable lengths for all strains collapsed when we divided the lengths of cables by the lengths of the cells in which they grew (Fig. 1, H and I). These results strongly suggest that cables grow to a length that matches cell length.

**Fig. 1.**
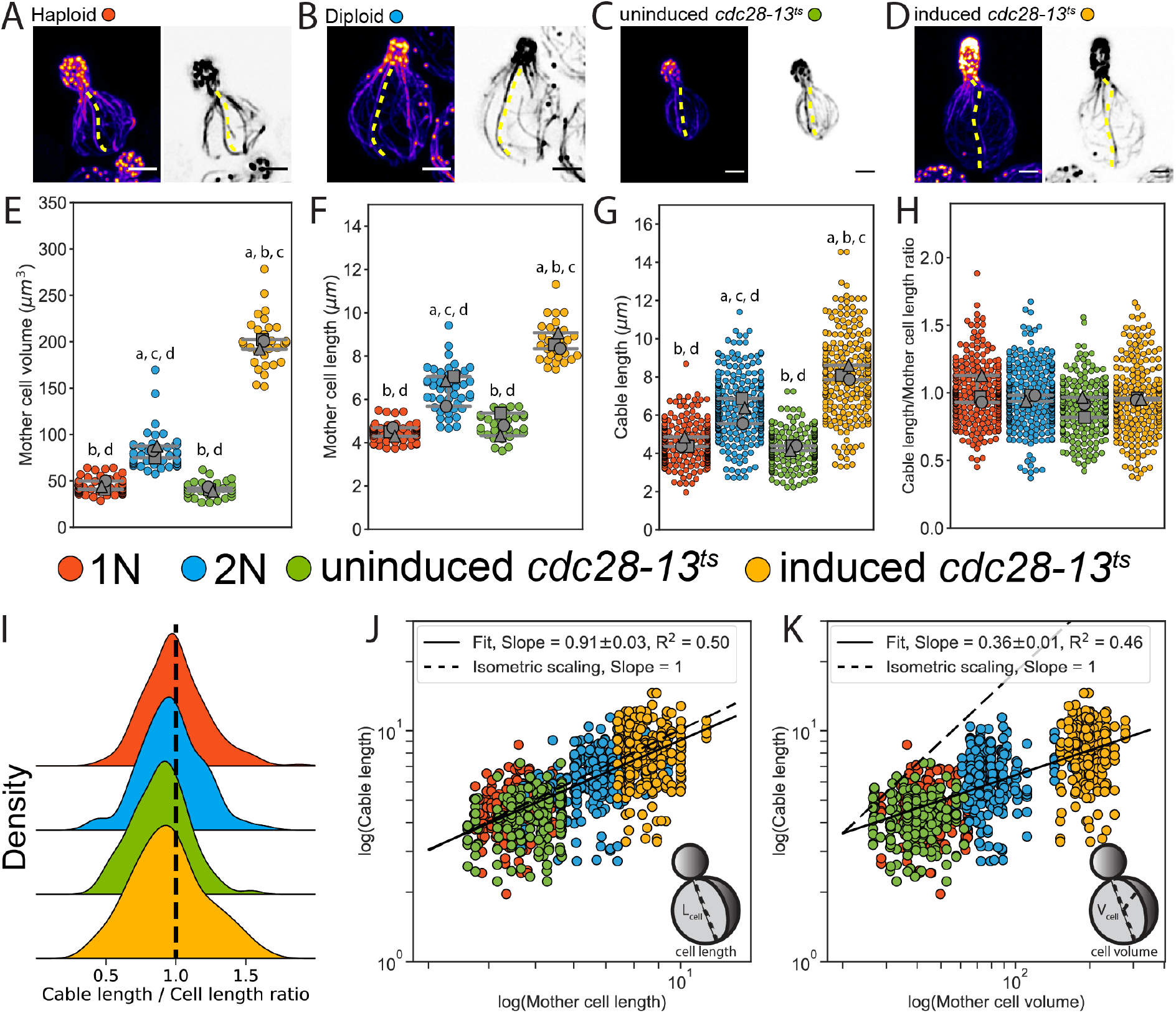
Actin cable length scales with cell length. **(A-D)** Representative images of haploid (A), diploid (B), uninduced *cdc28-13*^*ts*^ (C), and induced *cdc28-13*^*ts*^ (D) cells fixed and stained with labeled-phalloidin. Lengths of single actin cables are indicated (dashed lines) in maximum intensity projections (left, color) and single Z planes (right, inverted). Scale bar, 2 μm. **(E-F)** Mother cell volume (E) and length (F) measured in three independent experiments (≥30 cells/strain). Each data point is from an individual cell. Larger symbols represent the mean from each experiment. **(G-H)** Cable length (G) and ratio of cable length/cell length (H) measured from the same cells as in E and F (≥200 cables/strain). Each data point represents an individual cable. Larger symbols represent the mean from each experiment. Error bars, 95% confidence internals. Statistical significance determined by students t-test. Significant differences (p≤0.05) indicated for comparisons with haploid (‘a’), diploid (‘b’), uninduced *cdc28-13*^*ts*^ (‘c’), and induced *cdc28-13*^*ts*^ (‘d’). Complete statistical results in Table 2. **(I)** Probability density functions for ratios in H. **(J-K)** Cable lengths plotted against mother cell length (J) or volume (K) on double-logarithmic plots and fit using the power-law. Hypothetical isometric scaling (dashed line) is compared to experimentally measured scaling exponent (solid line).

Next, we used a power law analysis to rigorously test the scaling relationships of cable length with cell length and volume (Fig. 1, J and K). Generally, scaling relations can be described by the power law *y* = *A x*^*a*^, where *a* is the scaling exponent that reflects the relationship between the two measured quantities, *x* and *y* (*20*). This analysis revealed isometric scaling (*a*_*L*_ = 0.91 ± 0.03, *R*^2^ = 0.50) between cable length and cell length (Fig. 1J), whereas scaling between cable length and cell volume was hypoallometric (*a*_*V*_ = 0.36 ± 0.01, *R*^2^ = 0.46) (Fig. 1K). Therefore, in cells of vastly different size, the cable length directly scales with cell length, excluding the possibility that cable length scales with other dimensions, such as surface area or volume.

**Table 1:**
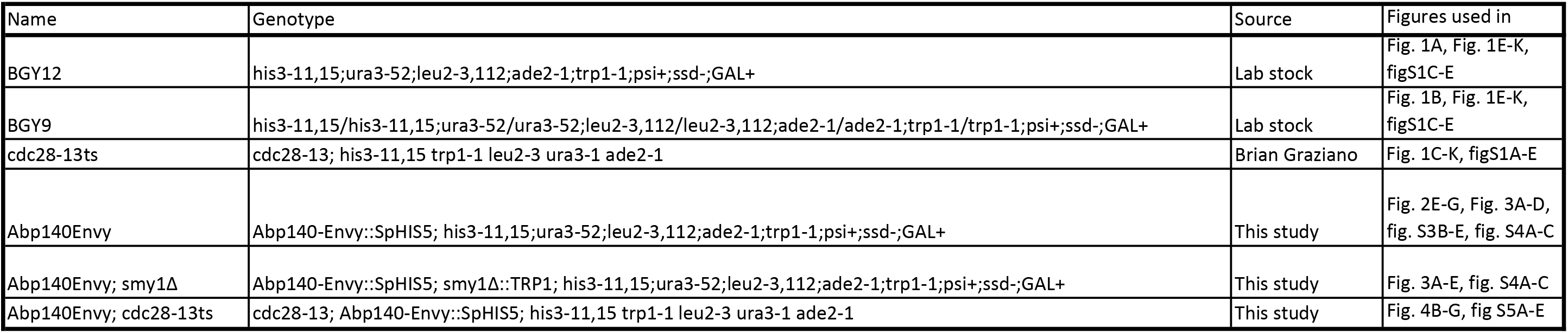
Yeast strains used in this study.

| Name | Genotype | Source | Figures used in |
| --- | --- | --- | --- |
| BGY12 | his3-11,15;ura3-52;leu2-3,112;ade2-1;trp1-1;psi+;ssd-;GAL+ | Lab stock | Fig. 1A, Fig. 1E-K, figS1C-E |
| BGY9 | his3-11,15/his3-11,15;ura3-52/ura3-52;leu2-3,112/leu2-3,112;ade2-1/ade2-1;trp1-1/trp1-1;psi+;ssd-;GAL+ | Lab stock | Fig. 1B, Fig. 1E-K, figS1C-E |
| cdc28-13ts | cdc28-13; his3-11,15 trp1-1 leu2-3 ura3-1 ade2-1 | Brian Graziano | Fig. 1C-K, figS1A-E |
| Abp140Envy | Abp140-Envy::SpHIS5; his3-11,15;ura3-52;leu2-3,112;ade2-1;trp1-1;psi+;ssd-;GAL+ | This study | Fig. 2E-G, Fig. 3A-D, fig. S3B-E, fig. S4A-C |
| Abp140Envy; smy1Δ | Abp140-Envy::SpHIS5; smy1Δ::TRP1; his3-11,15;ura3-52;leu2-3,112;ade2-1;trp1-1;psi+;ssd-;GAL+ | This study | Fig. 3A-E, fig. S4A-C |
| Abp140Envy; cdc28-13ts | cdc28-13; Abp140-Envy::SpHIS5; his3-11,15 trp1-1 leu2-3 ura3-1 ade2-1 | This study | Fig. 4B-G, fig S5A-E |

**Table 2:** Complete statistical results from students t-test conducted for data in Figure 1 and Supplemental Figure 1. P-values are indicated for each students t-test conducted.

| Figure 1E (Mother cell volume) | yBG12 | yBG9 | cdc28-13, uninduced | cdc28-13, induced |
| --- | --- | --- | --- | --- |
| yBG12 |  | 0.001 | 0.259 | 9.05E-07 |
| yBG9 | 0.001 |  | 4.45E-04 | 7.35E-06 |
| cdc28-13, uninduced | 0.259 | 4.45E-04 |  | 0.001 |
| cdc28-13, induced | 9.05E-07 | 7.35E-06 | 0.001 |  |

| Figure 1F (Mother cell length) | yBG12 | yBG9 | cdc28-13, uninduced | cdc28-13, induced |
| --- | --- | --- | --- | --- |
| yBG12 |  | 0.011 | 0.345 | 8.65E-05 |
| yBG9 | 0.011 |  | 0.035 | 0.010 |
| cdc28-13, uninduced | 0.345 | 0.035 |  | 0.001 |
| cdc28-13, induced | 8.65E-05 | 0.010 | 0.001 |  |

| Figure 1G (Cable length) | yBG12 | yBG9 | cdc28-13, uninduced | cdc28-13, induced |
| --- | --- | --- | --- | --- |
| yBG12 |  | 0.014 | 0.295 | 1.78E-04 |
| yBG9 | 0.014 |  | 0.008 | 0.012 |
| cdc28-13, uninduced | 0.295 | 0.008 |  | 7.40E-05 |
| cdc28-13, induced | 1.78E-04 | 0.012 | 7.40E-05 |  |

| Figure 1H (Cable length / Mother cell length) | yBG12 | yBG9 | cdc28-13, uninduced | cdc28-13, induced |
| --- | --- | --- | --- | --- |
| yBG12 |  | 0.528 | 0.218 | 0.40 |
| yBG9 | 0.528 |  | 0.22 | 0.408 |
| cdc28-13, uninduced | 0.218 | 0.22 |  | 0.30 |
| cdc28-13, induced | 0.40 | 0.408 | 0.30 |  |

| figure S1C (End-to-end distance) | yBG12 | yBG9 | cdc28-13, uninduced | cdc28-13, induced |
| --- | --- | --- | --- | --- |
| yBG12 |  | 0.014 | 0.392 | 1.79E-07 |
| yBG9 | 0.014 |  | 0.013 | 0.018 |
| cdc28-13, uninduced | 0.392 | 0.013 |  | 1.70E-06 |
| cdc28-13, induced | 1.79E-07 | 0.018 | 1.70E-06 |  |

| figure S1D (Tortuosity) | yBG12 | yBG9 | cdc28-13, uninduced | cdc28-13, induced |
| --- | --- | --- | --- | --- |
| yBG12 |  | 0.860 | 0.486 | 0.406 |
| yBG9 | 0.860 |  | 0.003 | 0.361 |
| cdc28-13, uninduced | 0.486 | 0.003 |  | 0.148 |
| cdc28-13, induced | 0.406 | 0.361 | 0.148 |  |

We considered two distinct models to explain the control of cable length. In both models, the length of a cable is determined by competing rates of actin assembly (*k*_+_) at the barbed ends of cables and disassembly (*k*_−_) at the pointed ends of cables (Fig. 2, A and B). Therefore, at any given time the extension rate of a cable is determined by the difference in its assembly and disassembly rates (Fig. 2B). In the boundary-sensing model, the assembly rate is greater than the disassembly rate until the extending cable physically encounters the rear of the cell, causing one or both rates to abruptly change (Fig. 2C, and fig. S2A)(*20*). This model predicts that the cable extension rate will be constant until the cable tip encounters the back of the cell. In contrast, the balance-point model requires that either the assembly rate, the disassembly rate, or both rates are length-dependent, and defines steady state cable length as the point at which these two rates are balanced (Fig. 2D, and fig. S2B)(*21*). In clear contrast to the boundary-sensing model, this model predicts that the cable extension rate will steadily decrease as the cable lengthens.

**Fig. 2.**
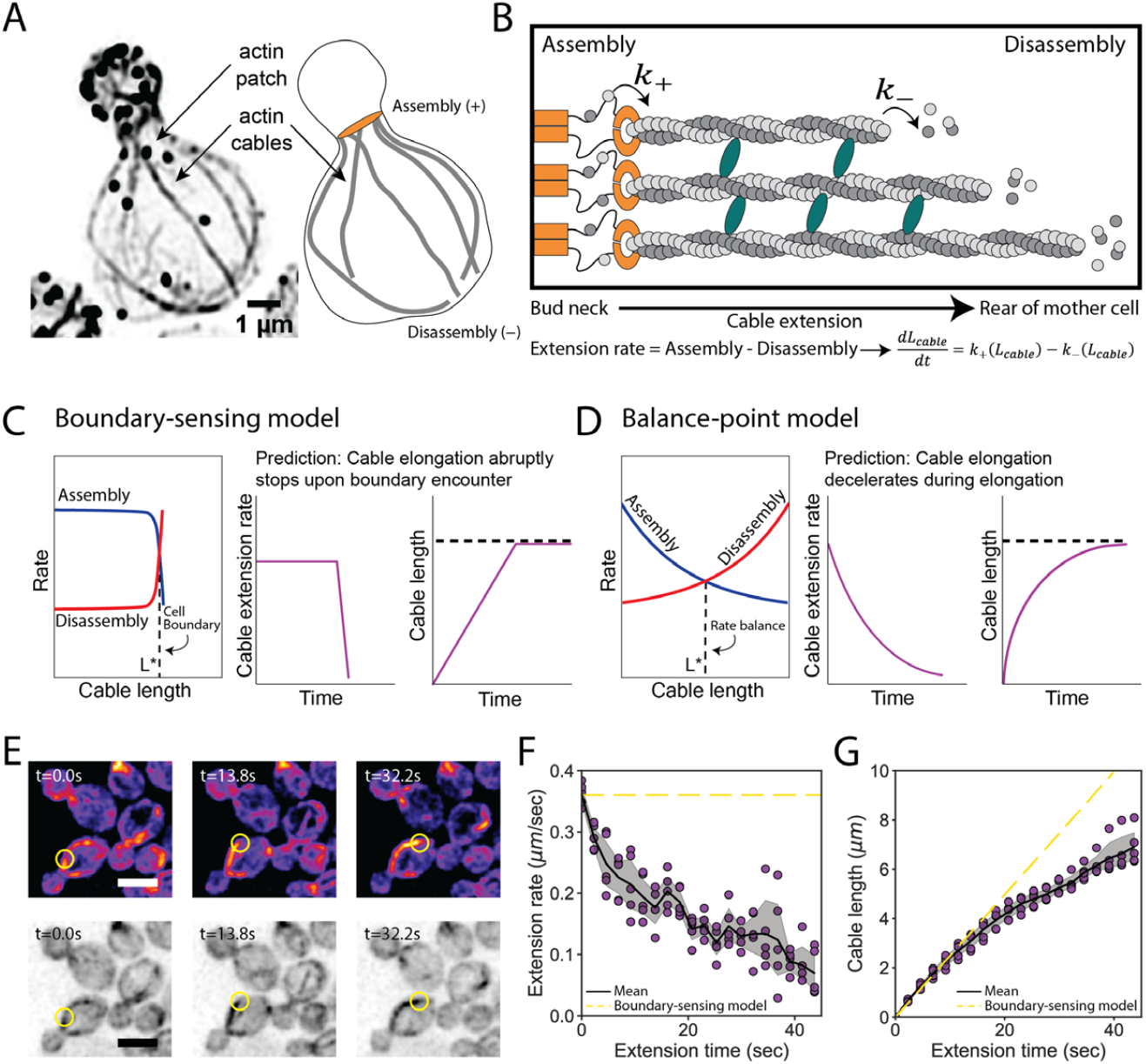
Models for control of actin cable length. **(A)** Actin staining in haploid cell (left) and cable traces (right). **(B)** Relevant parameters and equation for cable extension, where assembly (*k*_+_) and disassembly (*k*_−_) rates change as a function of cable length. Cables are polymerized by formins (orange) from actin monomers (gray), bundled by crosslinkers (blue), and disassembled by factors not shown. Cable extension rate is the difference in assembly and disassembly rates. **(C-D)** Two models for cable length control. Additional information in fig. S2. **(E)** Maximum intensity projection of haploid cell expressing cable marker (Abp140-GFP) shown in color (top panels) and inverted gray scale (bottom panels). Yellow circle highlights tip of elongating cable over time. Scale bar, 5 μm. **(F-G)** Extension rate (F) and length (G) measured in five independent experiments (n= 82 cables). Symbols at each time point represents mean for individual experiment. Solid lines and shading, mean and 95% confidence interval for all five experiments. Dashed yellow lines, predictions of boundary-sensing model in C.

To directly test the predictions of the two models, we used live imaging to track the tips of cables as they grew from the bud neck into the mother cell (Fig. 2E, and movie S1, and fig. S3, A, B, and C). Initially cables extended at 0.36 ± 0.02 μm s^−1^, and as they grew longer their extension rates steadily decreased (Fig. 2F, fig. S3D). Accordingly, we observed greater changes in cable length during earlier phases of cable growth (Fig. 2G, fig. S3E). Thus, as cables lengthen their growth rate decelerates.

Our experimental observations above support a balance-point model in which steady state cable length is reached when the assembly and disassembly rates are the same. In this model the rate of cable extension at any given time is given by the difference between the assembly and disassembly rates, which we call the feedback function, *f*. To account for the observed scaling of cable length with cell length (Fig. 1, H and I), we assume that *f* depends on the cable length (*L*_*cable*_) scaled by the cell length (*L*_*cell*_), i.e., *f*(*L*_*cable*_, *L*_*cell*_) = *f*(*L*_*cable*_/*L*_*cell*_). The steady state cable length 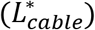 is reached when the feedback function equals zero, 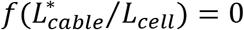. Therefore, the scale-invariant feedback function leads to the scaling of 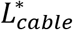 with *L*_*cell*_ seen in Fig. 1J. (Further mathematical details in methods.)

Smy1 is a factor implicated in cable length control, and therefore we considered whether it might be required for cable deceleration. It has been reported that cables are longer in *smy1*Δ compared to wildtype cells, and that Smy1 directly inhibits Bnr1-mediated actin assembly (*17*, *22*). Further, Smy1 is transported by myosin along cables to the bud neck where Bnr1 is anchored. Based on these observations, an ‘antenna mechanism’ has been proposed in which longer cables deliver more Smy1 to slow cable extension and limit cable length (*23*). We confirmed the increase in cable length in *smy1*Δ cells (Fig. 3A, and fig. S4A and B)(*17*), but found no difference in cable deceleration (Fig. 3, B and C). Instead, we observed an increase in the initial cable extension rate in *smy1Δ* (0.42 ± 0.04 μm s^−1^) compared to wildtype cells (0.35 ± 0.02 μm s^−1^) (Fig. 3, D and E). Thus, Smy1 influences cable length by limiting the initial cable growth rate (Fig. 3F), but does not provide the feedback that results in cable deceleration.

**Fig. 3.**
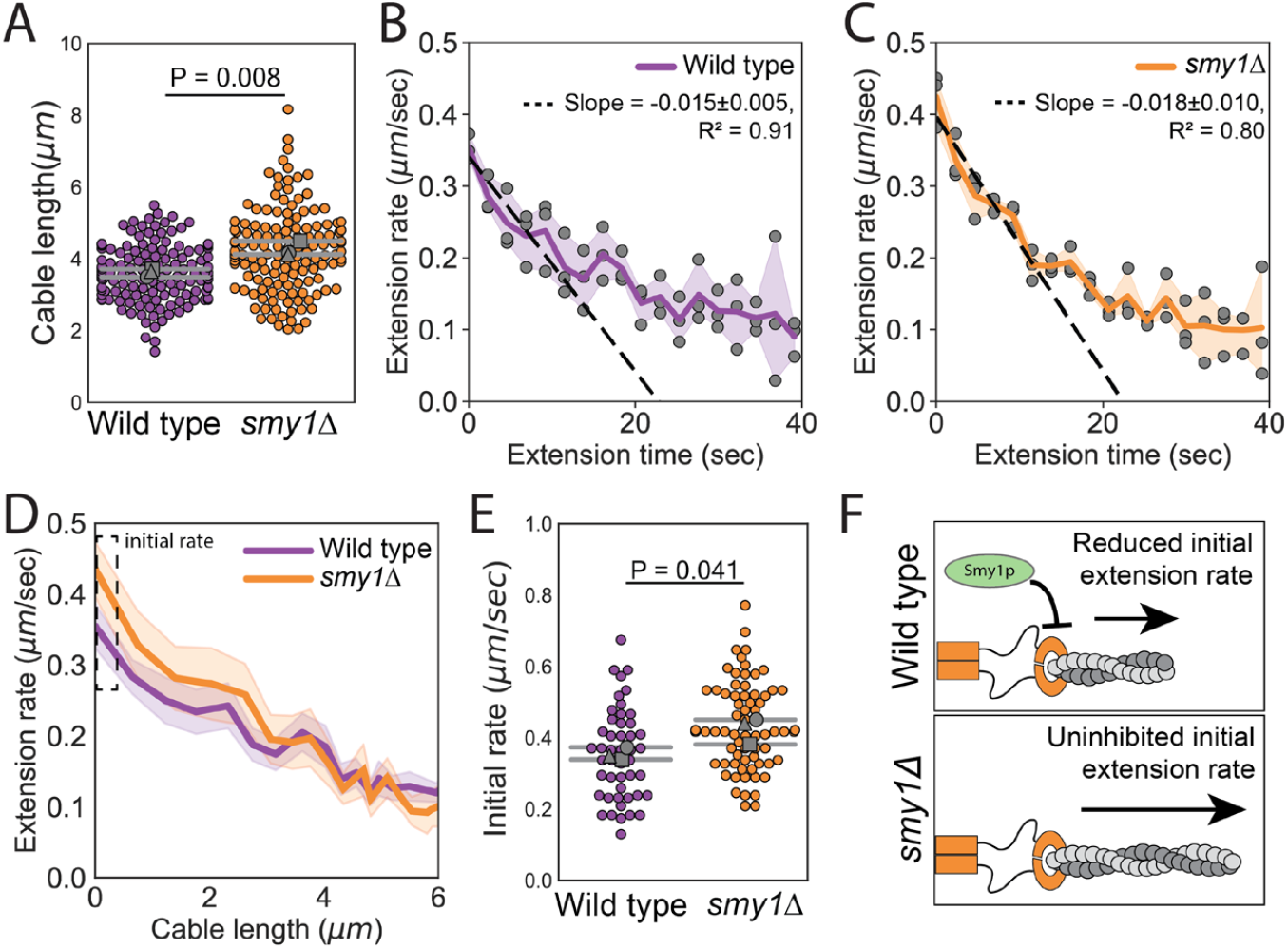
Smy1 controls initial cable extension rate. All data are from three independent experiments. **(A)** Cable lengths (≥130 cables/strain). Each data point represents an individual cable. Larger symbols, mean from each experiment. Error bars, 95% confidence internals. Statistical significance determined by students t-test. **(B-C)** Cable extension rates for wildtype (B) and *smy1*Δ (C) yeast (≥47 cables/strain). Symbols, mean from each experiment. Solid lines and shading, mean and 95% confidence interval for all experiments. Deceleration rates were derived from the slopes (±95%CI) of the dashed lines, which were determined by linear regression using the first ~10 seconds of extension. **(D)** Average extension rate as a function of cable length. Solid lines and shading, mean and 95% confidence interval for all experiments. Dashed box highlights region of no overlap in confidence intervals. **(E)** Initial cable extension rate for each strain. Small symbols, individual cables. Larger symbols, mean from each experiment. Error bars, 95% confidence intervals. Statistical significance determined by students t-test. **(F)** Cartoon comparing cable extension in wildtype and *smy1*Δ cells.

A key prediction of our balance-point model is that cable extension rates should depend on cell length, i.e., a cable of a given length should grow faster (or slow down more gradually) in longer cells compared to shorter cells (Fig. 4A, top). Further, it predicts that the cable extension rate profiles from cells of different lengths will collapse when cable length is normalized to cell length (Fig. 4A, bottom; predictions of model derived in methods). To test these predictions, we compared cable extension dynamics in uninduced and induced *cdc28-13*^*ts*^ cells (Fig. 4, B and C, fig. S5, A and B, and movies S2 and S3). When cables began to grow, they extended at similar rates in shorter and longer cells (fig. S5C). However, as the cables grew longer they decelerated more gradually in the longer cells, leading to longer cables (Fig. 4, D, E, and F, fig. S5D). Linear regression analysis further revealed that the magnitude of this difference in deceleration matches the predictions of our balance-point model (Fig. 4, D and E; details in methods). Additionally, cables extended with similar dynamics when cable length was normalized to cell length (Fig. 4G), as predicted by our model.

**Fig. 4.**
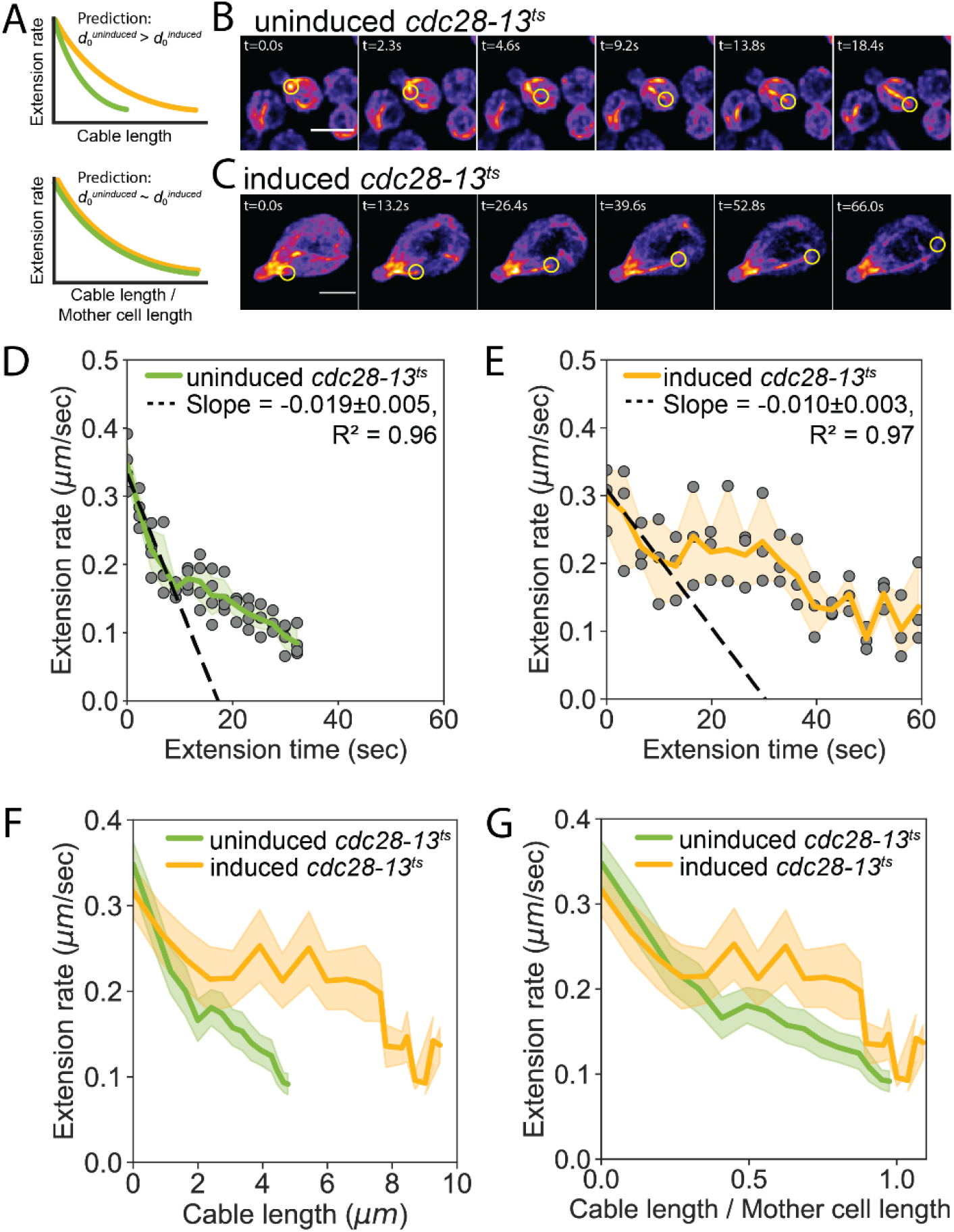
Cell length-dependent deceleration of actin cable growth. **(A)** Predictions of balance-point model comparing how cable deceleration (‘d’) changes as a function of cable length (top graph) in shorter (green curve) and longer (yellow curve) cells. This difference in the deceleration profiles is eliminated when cable length is normalized to cell length (bottom graph). **(B-C)** Maximum intensity projections of uninduced (B) and induced *cdc28-13*^*ts*^ (C) cells expressing cable marker (Abp140-GFP). Yellow circle highlights tip of elongating cable over time. Scale bar, 5μm. **(D-E)** Cable extension rates for uninduced (D) and induced *cdc28-13*^*ts*^ (E) cells, from at least three independent experiments (≥57 cables/strain). Symbols and shading, mean and 95% confidence intervals for all experiments. Deceleration rates were derived from the slopes (±95%CI) of the dashed lines, which were determined by linear regression using the first ~10 seconds of extension. **(F-G)** Average extension rates in uninduced and induced *cdc28-13*^*ts*^ cells (data from experiments in D and E) plotted as a function of cable length (F), or the ratio of cable length/cell length (G). Solid lines and shading, mean and 95% confidence interval for all experiments.

Collectively, our observations demonstrate that cables grow until their length matches the length of the cell, and that this is achieved by length-dependent cable deceleration. The precise mechanism providing the feedback to enable cable deceleration is not yet clear. However, it could be controlled by a gradient of actin disassembly-promoting activity that is highest at the rear of the cell. This would result in a greater disassembly rate for cables as they get longer and approach the rear of the cell. Further, such a gradient would be shallower in longer cells compared to shorter cells, accounting for the cell length-sensitive cable deceleration (fig. S2D). This mechanism also would allow cables to sense the rear of the cell without requiring physical interactions with that boundary. Alternatively, there could be length-dependent inhibition of cable assembly, e.g., conceptually similar to the antenna mechanism (*23*). However, because the site of cable assembly is fixed in cells (bud neck), whereas the site of cable disassembly changes as the cable lengthens, we favor a model involving length-sensitive tuning of the disassembly rate.

It has recently been shown for other subcellular structures (e.g., nucleus, spindle, centrosome, and nucleolus) that their sizes scale with cell volume, and this scaling is explained by limiting pool models (*3–5*, *8–10*). However, we found that polarized actin cables scale with cell length rather than volume. This length control cannot be explained by a limiting pool mechanism, and instead is explained, both theoretically and experimentally, by a balance-point model. These results reveal a new strategy by which cells solve engineering challenges, enabling them to scale internal structures with the linear dimensions of the cell (*24*). Similar principles may underlie the length control of other polarized, linear actin structures, such as filopodia and stereocilia. Further, related strategies may be used to control the growth of radial microtubule arrays that reach the cell periphery (*25*, *26*), and may explain the scaling relationships observed between flagellar length and cell length (*27*) and between contractile ring diameter and cell diameter (*28*).

## Acknowledgments

We thank Ariel Amir, Lishibanya Mohapatra, James Moseley, Rob Phillips, Aldric Rosario, and Alison Wirshing for thoughtful discussions on cable length control and comments on the manuscript. We are also grateful to Brian Graziano for sharing the *cdc28-13*^*ts*^ strain.

## Funding

This research was supported by the Brandeis NSF Materials Research Science and Engineering Center grant (MRSEC-142038), an award from the NSF Postdoctoral Research Fellowships in Biology Program to S.G.M. (Grant No. 2010766), grants from NSF (DMR-1610737) and the Simons Foundation (www.simonsfoundation.org/) to J.K., and a grant from the NIH to B.L.G. (R35 GM134895).

## Author contributions

SGM, JK, and BLG designed the experiments and wrote the manuscript. SGM performed experiments and analyzed the data.

## Competing interests

Authors declare no competing interests.

## Data and materials availability

Data are available in the main text or in the supplementary material. All images are archived at Zenodo. Source code is available on GitHub: https://github.com/shanemc11/Yeast_actin_cables.

## Supplementary Materials

Materials and Methods

Figure legends

Figures S1–S5

Tables S1-S2

Captions for Movies S1 to S3

Movies S1-S3

## Materials and Methods

### Plasmids and yeast strains

All strains (see Supplementary Table 1) were constructed using standard methods. To integrate a bright GFP variant (Envy) at the C-terminus of the endogenous *ABP140* gene, primers were designed with complementarity to the 3’ end of the GFP cassette and the C-terminal coding region of *ABP140.* PCR was used to generate amplicons from the pFA6a-GFP-His3MX template that allow for selection of transformants using media lacking histidine. The parent strains, BGY12 (haploid) and *cdc28-13*^*ts*^, were transformed with PCR products, and transformants were selected by growth on synthetic media lacking histidine. Similarly, *smy1Δ* strains were generated by replacement of *SMY1* with the *TRP1* auxotrophic marker by designing primers with complementarity to regions 40 base-pairs immediately up-stream and down-stream of the *SMY1* coding region. Deletion of *SMY1* was confirmed by genomic PCR with primers specific to the *TRP1* promoter and the 5’UTR region of *SMY1*. The *cdc28-13*^*ts*^ strain was a generous gift from Dr. Brian Graziano (UCSF).

### Induction of cell size changes

To induce enlargement of mother cells, *cdc28-13* cells were grown at the permissive temperature (25°C) overnight in synthetic complete media (SCM), then diluted to OD_600_ ~ 0.1 in fresh SCM. Cultures were then shifted to the restrictive temperature (37°C) for 8 hours (except for the experiments in figure S1A and B, where cultures were also shifted for only 4 hours). After this induction, cells were returned to the permissive temperature (25°C) for one hour of growth to allow cell polarization and bud growth, and then fixed or mounted for live-cell imaging.

### Quantitative analysis of actin cable length and architecture in fixed cells

Strains were grown at 25°C to mid-log phase (OD_600_ ~ 0.3) in yeast extract/peptone/dextrose (YEPD), or were first induced for cell size changes as indicated above. Then cells were fixed in 4.4% formaldehyde for 45 minutes, washed three times in phosphate-buffered saline (PBS), and stained with Alexa Fluor 488-phalloidin or Alexa Fluor 568-phalloidin (Life Technologies, Grand Island, NY) for ≥24 hours at 4°C. Next, cells were washed three times in PBS and imaged in mounting media (20mM Tris, pH 8.0, 90% glycerol). 3D stacks were collected at 0.22 μm intervals on a Zeiss LSM 880 using Airyscan super-resolution imaging equipped with 63× 1.4 Plan-Apochromat Oil objective lens. 3D stacks were acquired for the entire height of the cell. Airyscan image processing was performed using Zen Black software (Carl Zeiss). ImageJ was used to generate inverted greyscale and maximum projection images for analysis. For length analysis, we included every discernable cable in the cell that extended from the bud neck to some endpoint in the mother cell; the only cables excluded were the minority that became closely intertwined with other cables making it impossible to resolve their individual lengths. Using ImageJ, each individual cable was manually traced through the 3D stack, from the bud neck to their terminus in the mother cell, and then the xy-coordinates for each cable trace were exported into custom written Python scripts to compute cable length. Cell length was determined by measuring the distance from the bud neck to the distal end of the mother cell. Cell width was determined by measuring the widest point perpendicular to the cell length axis. Cell height was determined from the number of slices in the 3D stack and the interval size between slices. These values were recorded and imported into custom Python scripts to compute the ratio of cable length to mother cell length, the cell volume (using the ellipsoid formula), and to fit the scaling exponent for cable length versus mother cell length, width, and volume.

### Live-cell imaging and quantitative analysis of actin cable extension rate

Strains were grown at 25°C to mid-log phase (OD_600_ ~ 0.3) in either YEPD, or were first induced for cell size changes as indicated above, then harvested by centrifugation (30 seconds, 9000 x g). Media was decanted and cells were resuspended in 50 μL fresh media. Cells (~5 μL) were mounted onto 1.2% agarose pads made with SCM, and images were acquired on a Nikon i-E upright confocal microscope equipped with a CSU-W1 spinning-disk head (Yokogawa, Tokyo, Japan) and an Andor Ixon 897 Ultra CCD camera controlled by Nikon NIS-Elements Advanced Research software using a 100x, 1.45 NA objective. 3D stacks were acquired at 0.3 μm intervals for approximately half of the cell height with no time delay for 2 minutes (approximately 0.30-0.43 frames per second). Images were processed in ImageJ by generating maximum intensity projections of each stack and applying a Gaussian blur (sigma = 1) to facilitate manual tracking of cable tips, as they emerged from the bud neck and until they stopped extending. Individual cable trajectories were imported into custom Python scripts to compute the distance the cable tip travelled between each frame, the rate of extension between each frame, and the total distance travelled. The boundary-sensing model prediction depicted in figure 2F was determined by plotting the mean initial cable extension rate as a function of time. The boundary-sensing model prediction depicted in figure 2G was determined by using linear regression to measure the slope from the first ~10 seconds of cable extension. Initial cable extension rates (Fig. 3C and fig. S5A) were determined by computing the extension rate measured during the first time interval.

### Mathematics of the balance point model

The rate of change of the cable length with time is given by the difference between the assembly (*k*_+_) and disassembly (*k*_−_) rates,

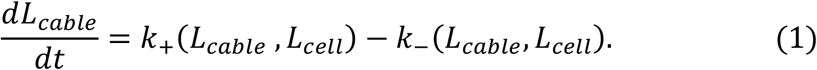

where we have made explicit the possibility that one or both rates depend on the length of the extending cable (*L*_*cable*_) and the cell length (*L*_*cell*_). The steady state length 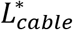 is the cable length at which the assembly and disassembly rates are the same.

To account for the scaling of the steady state length with the cell length (as observed in Fig. 1 H, I, and J), we make an additional assumption, namely that the feedback function, *f* ≡ *k*_+_ − *k*_−_, which determines the rate of cable extension, is a function of the ratio of the cable length to the cell length, i.e., *f*(*L*_*cable*_, *L*_*cell*_) = *f*(*L*_*cable*_/*L*_*cell*_). Thus our mathematical model of cable length control is described by the equation:

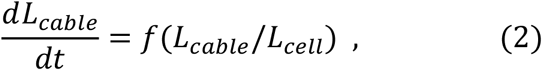

which is graphically summarized in supplemental figure 2C.

At the molecular scale, this feedback could be accomplished with a constant rate of cable assembly and a variable rate of disassembly controlled by a gradient of depolymerizing activity that is highest at the back of the cell; see supplemental figure 2D. In this mechanism, as the cable lengthens its distal end is subject to increasingly stronger depolymerizing activity. Further, the profile or decay-length of the gradient will scale with cell length. Such scaling of a cellular gradient with the linear distance between the two poles of the cell has been observed for the protein Bicoid in different size embryos, from different species of flies (*29*). Other experimental observations and theoretical models of such scale-invariant gradients are reviewed in (*30*).

Figures 4F and G are a direct test of our model. In figure 4F, we observe that the cable extension rate is dependent on cell length, consistent with equation 1. In figure 4G, we see that the two feedback functions, from cells of different size, collapse to a single function when the cable lengths are scaled by the cell length.

The scaling property of the feedback function immediately leads to scaling of steady state cable length with cell length. Namely, in steady state, the right-hand side of equation 1 is zero, which implies 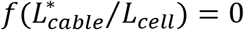. If the zero of the feedback function is *x*\* (i.e., *f*(*x*\*) = 0), then the steady state length 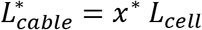, which is the scaling relation we observe in Fig. 1 H and J between the steady state length and the cell length.

The scaling property of the feedback function also makes a prediction for the initial rate of cable extension in cells of different size. Namely, for small cable lengths, when *L*_*cable*_ ≪ *L*_*cell*_, we can expand equation 2 into a Taylor series

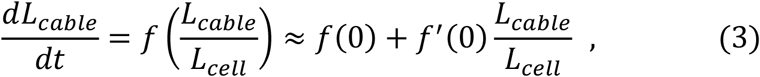

which states that the initial cable extension decreases linearly with the cable length (since *f*′(0) is negative) that is inversely proportional to cell length, *L*_*cell*_.

Equation 3, with the initial condition *L*_*cable*_(*t* = 0) = 0, can be solved for the cable length as a function of time,

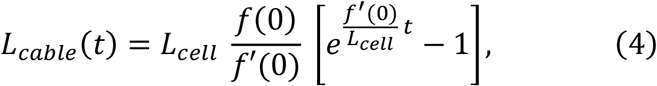

which in turn yields, by differentiation, an exponentially decreasing in time extension rate:

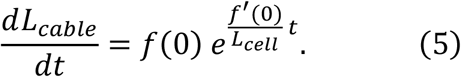

Since equations 4 and 5 only hold at early times when the cable length is much smaller than the cell length (roughly, first 10 seconds of cable extension; see figure 2G), we can further simplify equation 5 by expanding it into a Taylor series:

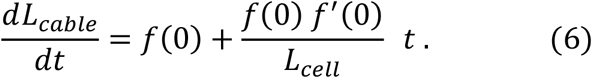

Equation 6 makes very specific predictions about the initial deceleration of cable extension, in particular our model (equation 2) predicts that the initial deceleration

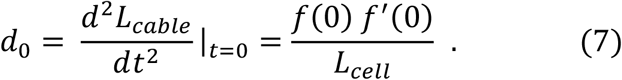

scales inversely with the cell length, which we observe in our experiments. Indeed, when comparing the initial deceleration of cable extension in the uninduced and induced *cdc28-13*^*ts*^ mutant cells in Fig. 4 D and E, we find 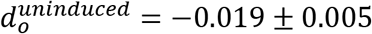 and 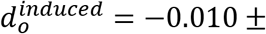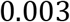, respectively. The ratio of the deceleration in the small and large cells, 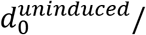 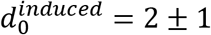, is consistent with our prediction that this ratio should equal the inverse of the ratio of the cell lengths, 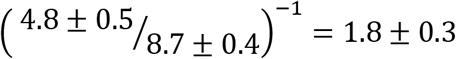.

Our model also makes an interesting quantitative prediction for cable extension in *smy1*Δ cells. As described in the main text, in these mutant cells the initial cable extension rate is higher than in wildtype cells, *f*(0)^*wt*^ = 0.35 ± 0.02 *μm*/*s*, versus *f*(0)^Δ*smy*1^ = 0.42 ± 0.04 *μm*/*s*. Equation 7 says that this difference in initial extension rate will lead to a proportional increase in the initial deceleration. Indeed, linear fits to the cable extension rate, as a function of time over the first 10 s, yield, 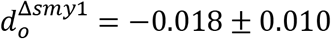 and 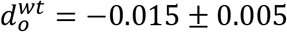, for *smy1*Δ and wild type cells, respectively. The ratio of these two, 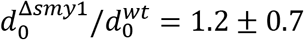, matches the ratio of the initial extension rates, *f*(0)^*Δsmy*1^/*f*(0)^*wt*^ = 1.2 ± 0.2, as predicted by our equation.

Finally, our model also makes a qualitative prediction about the probability distribution of cable lengths at steady state. Namely, the feedback function near the steady state cable length, 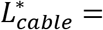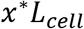 can be Taylor expanded to

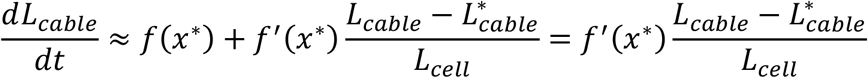

which shows that the strength of the feedback function (*f*) diminishes with cell length. This in turn implies that the steady state fluctuations of cable length will be larger in longer cells, which is consistent with data in figure 1G. It is important to note that the above arguments pertain to cable length fluctuations over time, whereas the data in figure 1G show cell-to-cell fluctuations in cable length, which could be influenced by cell-to-cell heterogeneity in some of the factors that affect cable assembly. Further experiments that carefully delineate between different sources of cable length fluctuations could provide more detailed tests of our model.

---

**Fig. S1.**
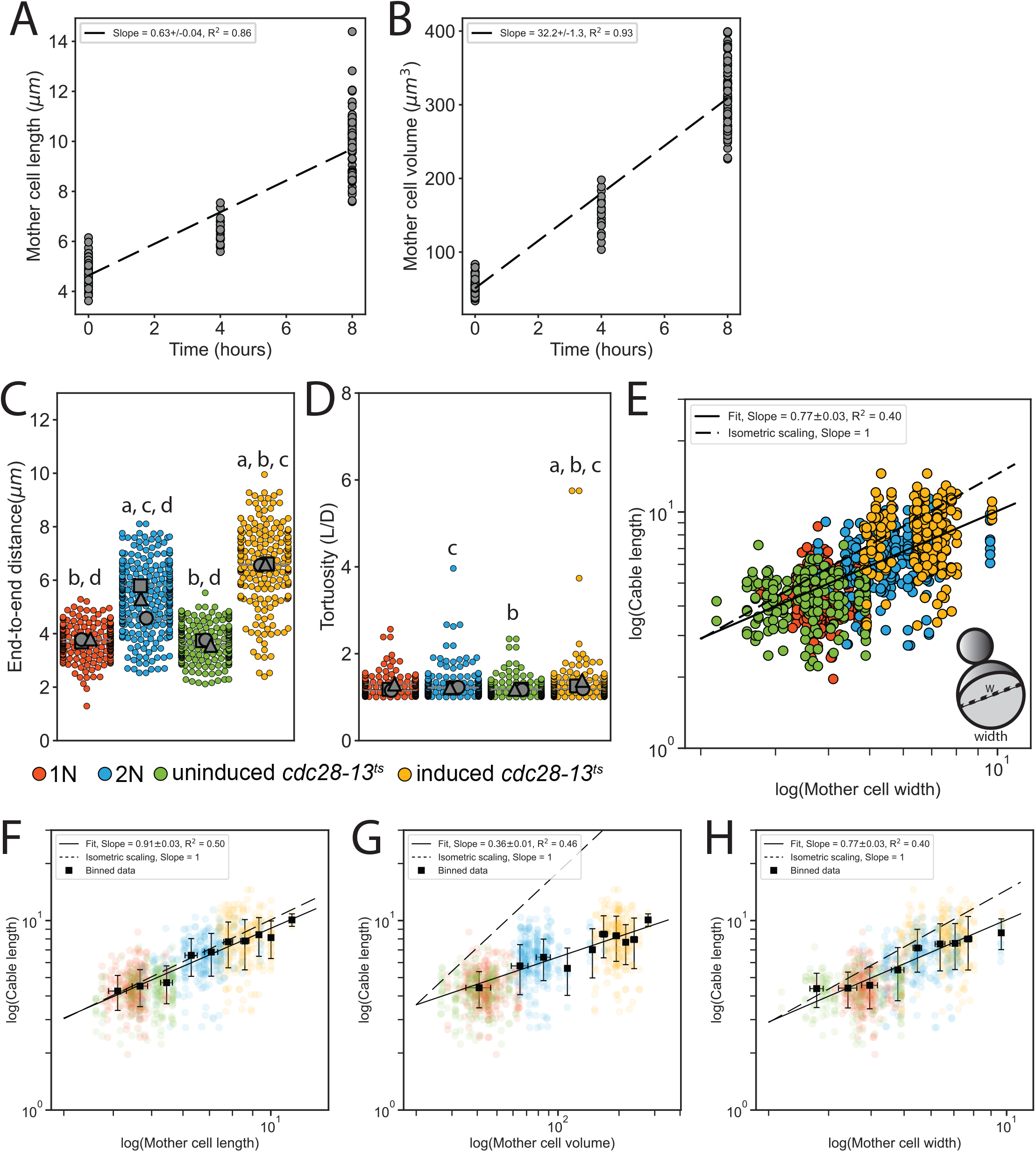
Changes to actin cable architecture in cells of different size. **(A-B)** Changes in mother cell length (A) and cell volume (B) of *cdc28-13*^*ts*^ cells after growth at the restrictive temperature for the indicated times (≥20 cells/strain). Dashed line indicates results from linear regression. **(C-D)** End-to-end cable distance (C) and length/distance, or tortuosity (D) measured in three independent experiments (≥200 cables/strain). Each data point represents an individual cable. Larger symbols, mean from each experiment. Error bars, 95% confidence internals. Statistical significance determined by students t-test. Significant differences (p≤0.05) indicated for comparisons with haploid (‘a’), diploid (‘b’), uninduced *cdc28-13*^*ts*^ (‘c’), and induced *cdc28-13*^*ts*^ (‘d’). Complete statistical results in Table 2. **(E)** Cable lengths plotted against mother width on a double-logarithmic plot, and fit using the power-law. This analysis indicated that scaling between cable length and cell width is hypoallometric (*a*_*W*_ = 0.77 ± 0.03, *R*^2^ = 0.40), in contrast with the observed isometric scaling between cell length and cable length (Fig. 1J) Hypothetical isometric scaling (dashed line) is compared to experimentally-measured scaling exponent (solid line). **(F-H)** Binned data (black squares, mean ± standard deviation) from figures 1J, 1K and supplemental figure 1E.

**Fig. S2.**
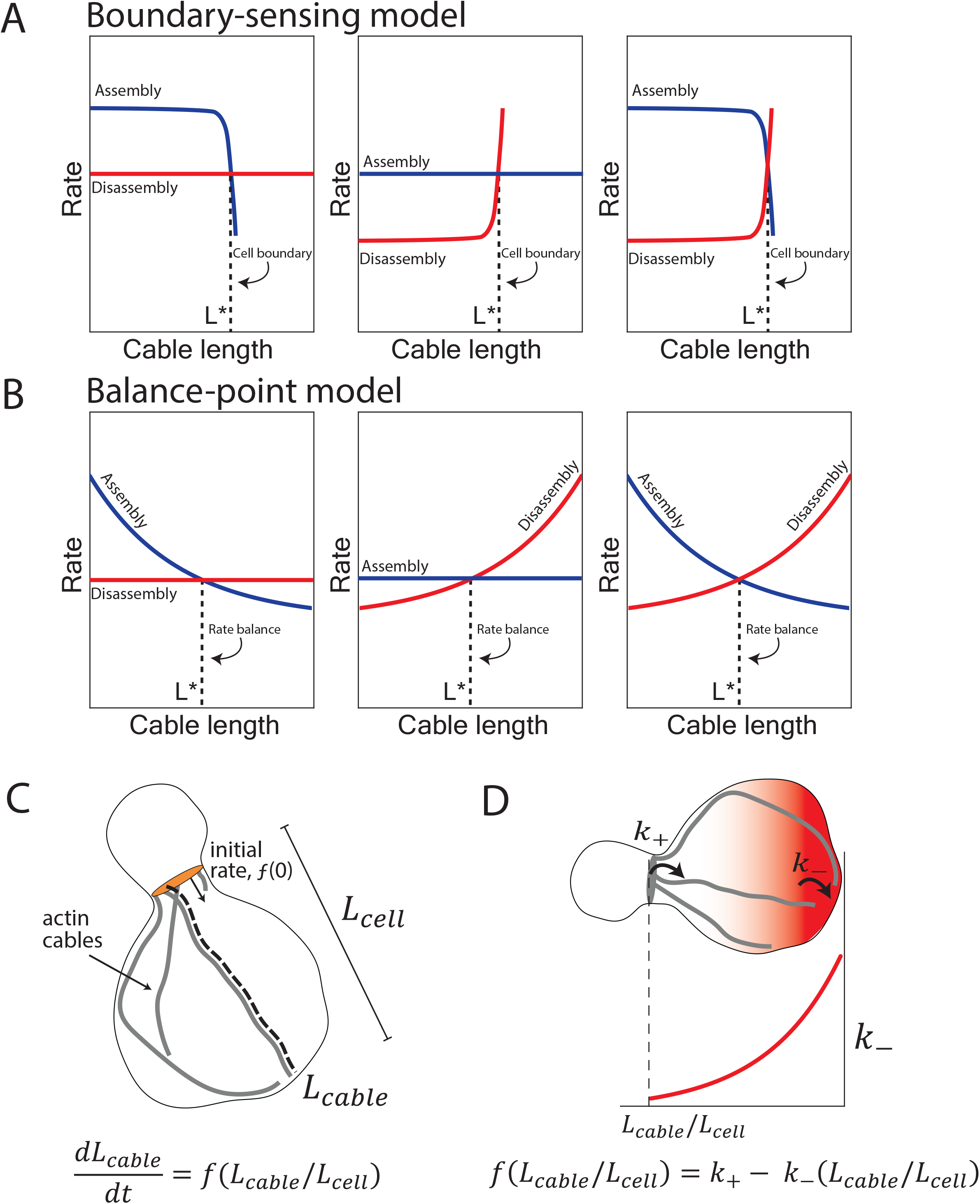
Boundary-sensing and balance-point models of length regulation. **(A)** The boundary-sensing model proposes that cable length scales with mother cell length due to the growing tip of the cable physically encountering the rear of the mother cell (cell boundary). In this model, cable assembly and disassembly rates are length-independent until the boundary is reached. As the cable is growing (extending), the assembly rate must be greater than the disassembly rate. When the cable encounters the boundary, there is an abrupt shift in rates to prevent further growth of the cable: the assembly rate rapidly decreases (left panel), the disassembly rate rapidly increases (middle panel), or both rates change (right panel). Based on these changes to the assembly and/or disassembly rates, the boundary sensing model predicts that the rate of cable extension remains constant until the cable physically encounters the rear of the mother cell, and then it abruptly decreases (Fig. 2C). This behavior results in a cable that grows linearly with time until it reaches the boundary (Fig. 2C). **(B)** In contrast, in balance-point model, the assembly rate (left panel), the disassembly rate (middle panel), or both rates (right panel) are length-dependent, and the intersection of these two rates (rate balance) produces a steady-state cable length (dashed line). Therefore, the balance-point model predicts that the cable extension rate will decelerate as a cable grows longer (Fig. 2D). In the balance-point model, changes in cable length are expected to be greater during initial time points in a cable’s growth, and gradually decline until the steady-state length is reached (Fig. 2D, dashed line). **(C)** Cartoon of yeast cell with actin cables (gray) extending from formins (orange) at the bud neck. Relevant parameters and mathematical form for the derived balance-point model are indicated, where the change in cable length as a function of time is controlled by the feedback function, *f* (additional details in methods). **(D)** Cartoon depiction of a mechanism where actin cable length is scaled with cell length by a constant rate of cable assembly (*k*_+_), and a variable rate of disassembly (*k*_−_) controlled by a gradient of depolymerizing activity (red shading). The gradient is highest at the back of the mother cell, which leads to the disassembly rate (red line, lower plot) increasing as the cable lengthens. By this mechanism, the cable length will scale with cell length.

**Fig. S3.**
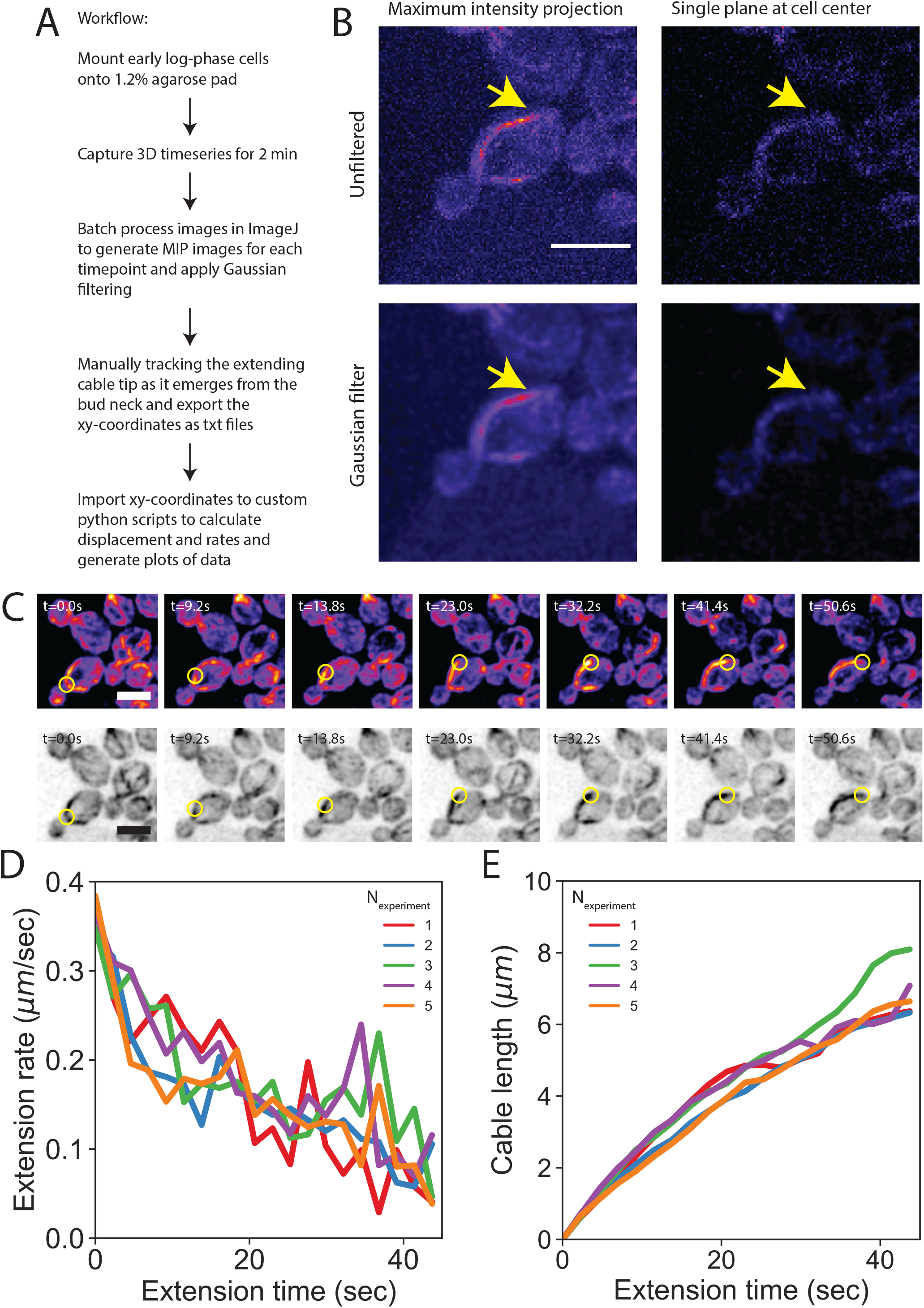
3D timeseries imaging workflow. **(A**) Yeast cells grown to early log-phase were mounted onto 1.2% agarose pads made with synthetic complete media and imaged on a spinning disk confocal microscope. 3D stacks were acquired at 0.3 μm intervals for approximately half of the cell height with no time delay for 2 minutes (approximately 0.30-0.43 frames per second). Images were processed using custom ImageJ macros to generate maximum intensity projections for each stack and apply a Gaussian blur (sigma = 1) to facilitate manual tracking of cable tips from the point where they emerged from the bud neck until they ceased extending. Individual trajectories were analyzed using custom Python scripts to compute the distance of cable tip extension between each frame, the rate of cable extension between each frame, and the total extension distance starting from initial growth at the bud neck. **(B)** Tracking the entire lifetime of a single cable was made possible by capturing 3D timeseries images in conjunction with our image analysis pipeline. Representative images of a single time point of a cell expressing the cable marker Abp140-GFP (integrated) show that 2D timeseries images do not have sufficient signal-to-noise ratio for long term cable tracking (upper right panel). Application of a Gaussian blur to the timeseries images helps enable reliable tracking of cables (bottom panels). Scale bar, 5 μm. **(C)** Maximum intensity projection of haploid cells expressing a cable marker (Abp140-GFP) shown in color (top panels) and inverted gray scale (bottom panels). Yellow circle highlights tip of an elongating cable over time. Scale bar, 5 μm. **(D-E)** Mean cable extension rates (D) and mean cable lengths (E) for the five independent experiments shown in figure 2F and 2G (n= 82 cables). Line color indicates the mean of each replicate.

**Fig. S4.**
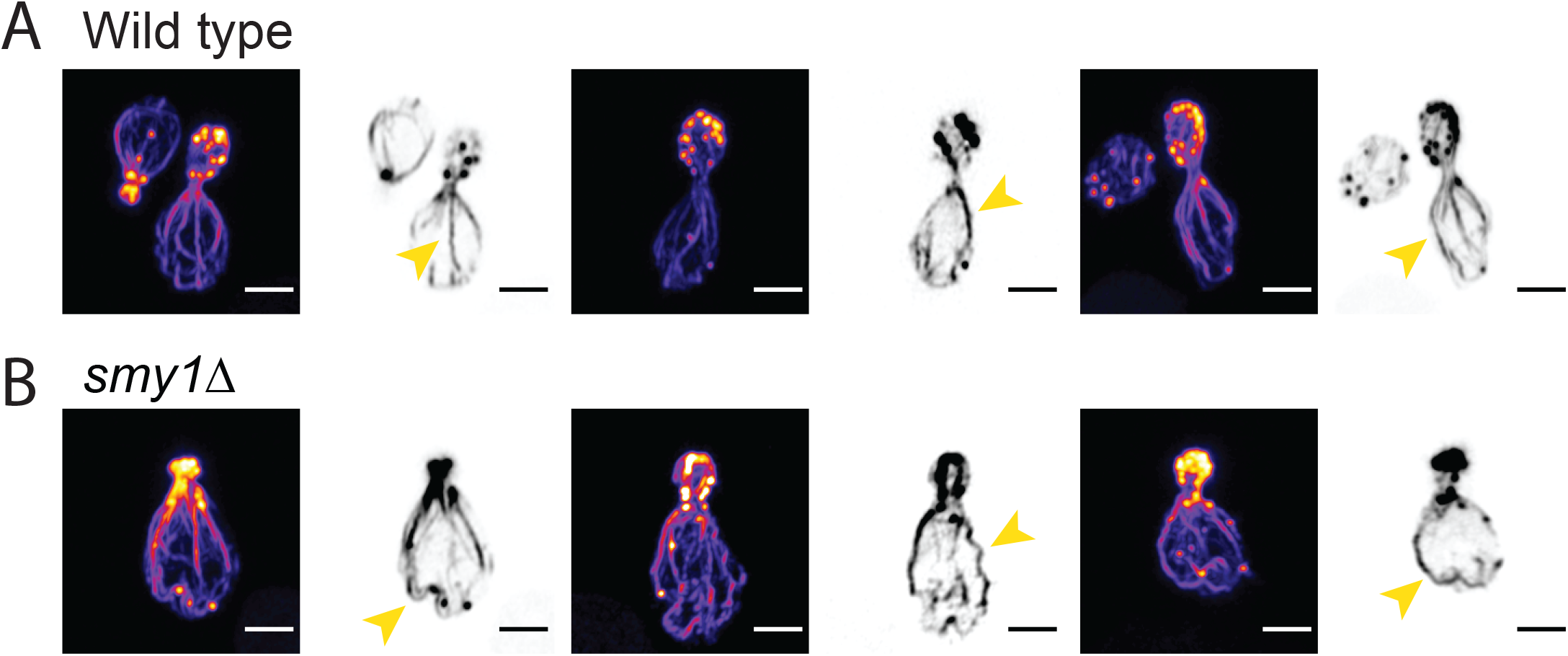
Altered actin cable length and architecture in *smy1Δ* cells. **(A-B)** Representative images of fixed haploid wildtype (A) and *smy1Δ* (B) cells stained with fluorescent phalloidin to label F-actin. Actin cables in wildtype cells grow to reach the back of the mother cell and have a relatively straight appearance (yellow arrows in A), whereas cables in *smy1Δ* are abnormally long and have a ‘wavy’ appearance (yellow arrows in B), in agreement with previous studies (*17*, *22*). Scale bar, 2 μm.

**Fig. S5.**
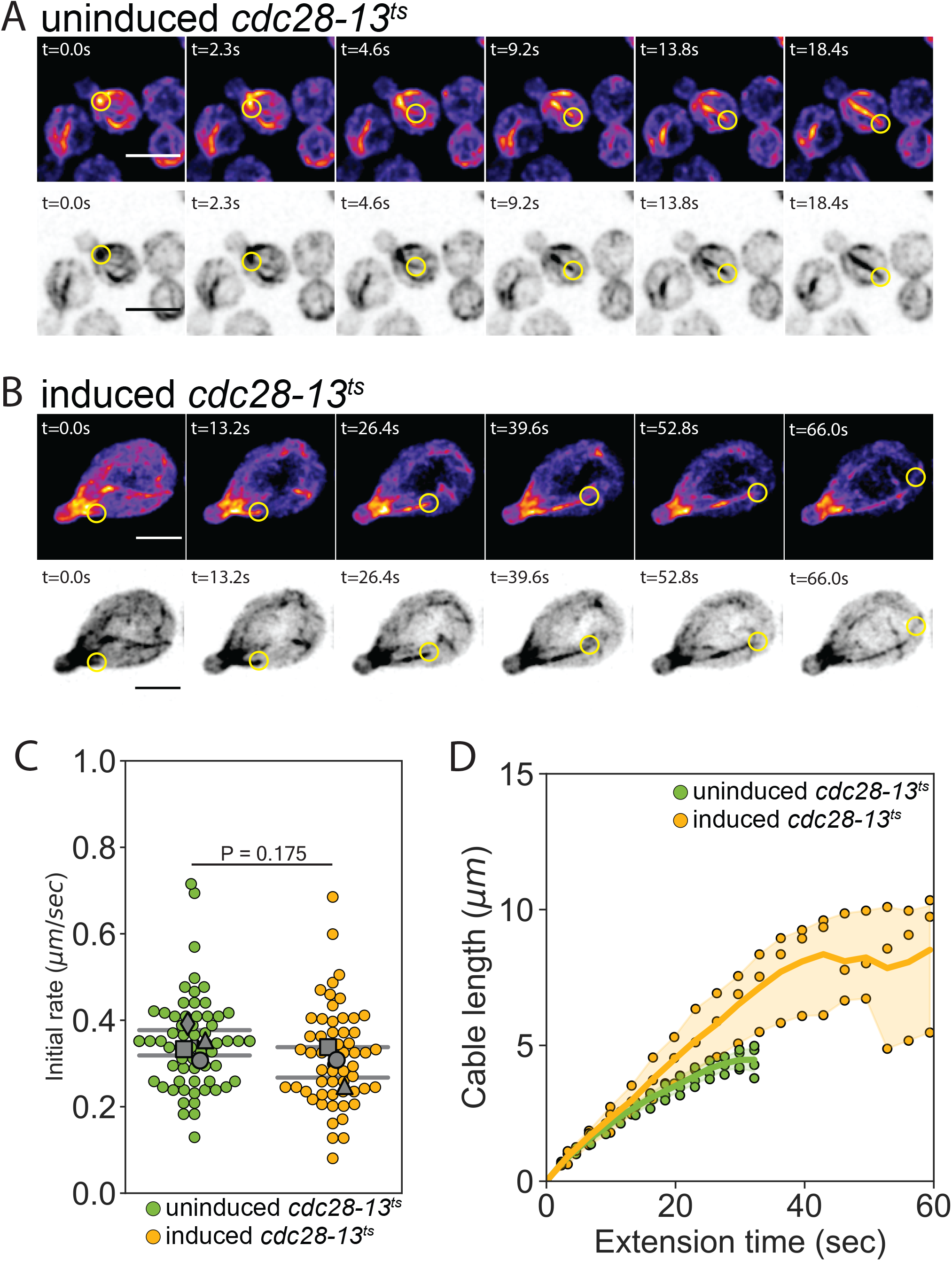
Cable extension dynamics are cell-length dependent. **(A-B)** Maximum intensity projections of uninduced (A) and induced *cdc28-13*^*ts*^ (B) cells expressing cable marker (Abp140-GFP) shown in color (top panels) and inverted gray scale (bottom panels). Yellow circle highlights tip of elongating cable over time. Scale bar, 5μm. **(C)** Initial cable extension rate for uninduced (green) and induced *cdc28-13*^*ts*^ (yellow) cells. Data are from live-cell imaging experiments as in figure 4. Small symbols, individual cables. Larger symbols, mean from at least three independent experiments (≥57 cables/strain). Error bars, 95% confidence intervals. Statistical significance determined by students t-test. **(D)** Analysis of data from C, showing changes in actin cable length (D) plotted versus extension time. Symbols, mean from each experiment. Solid lines and shading, mean and 95% confidence interval for all experiments.

Table S1. **Yeast strains used in this study.**

The genotype, source, and related data are indicated for each strain used in this study.

Table S2. **Complete results from statistical tests performed in this study.**

The p-value is indicated for student’s t-test used to compare the data presented in Figure 1 and Supplemental Figure 1.

**Movie S1.**

Maximum intensity projection of haploid cells expressing a cable marker (Abp140-GFP) shown in color. Yellow circle highlights tip of an elongating cable over time. Video is played at 10 frames per second and time (seconds) is indicated in the top left corner. Scale bar, 5 μm.

**Movie S2.**

Maximum intensity projections of uninduced *cdc28-13*^*ts*^ cells expressing cable marker (Abp140-GFP) shown in color (top panels). Yellow circle highlights tip of elongating cable over time. Video is played at 10 frames per second and time (seconds) is indicated in the top left corner. Scale bar, 5μm.

**Movie S3.**

Maximum intensity projections of induced *cdc28-13*^*ts*^ cells expressing cable marker (Abp140-GFP) shown in color (top panels). Yellow circle highlights tip of elongating cable over time. Video is played at 10 frames per second and time (seconds) is indicated in the top left corner. Scale bar, 5μm.

## Notes

### Competing Interest Statement

The authors have declared no competing interest.

